# The nucleus of the lateral olfactory tract is required for learning odor-guided food avoidance

**DOI:** 10.64898/2026.07.16.738920

**Authors:** Naheel Lawabny, Aya Dhamshy, Tirzah Kreisel, Tamar Licht, Dan Rokni

**Affiliations:** Department of Medical Neurobiology, Faculty of Medicine and IMRIC, The Hebrew University of Jerusalem, 9112102 Jerusalem, Israel; The Edmond and Lily Safra Center for Brain Sciences, The Hebrew University of Jerusalem, 9190401 Jerusalem, Israel

## Abstract

Foraging animals must continually evaluate whether potential food sources are safe to consume. Such decisions rely on both innate preferences and learned associations formed through prior experience. Consumption of a food followed by malaise or sickness leads to avoidance of that food, reflecting learned associations between sensory cues that identify a food and its negative post-ingestive consequences. Although this form of learning has been studied extensively using taste cues, taste signals arise only after oral contact, at which point exposure to potential toxins has already occurred. Olfactory cues, in contrast, provide information about potential food sources prior to consumption, yet how odors acquire aversive value remains unclear. Here, using targeted chemogenetic perturbations, we identify the nucleus of the lateral olfactory tract (NLOT), which has direct bidirectional connectivity to olfactory regions and the basolateral amygdala, as a critical circuit element for odor-aversion learning. We find that the NLOT is required for conditioned odor aversion but is dispensable for other odor-guided behaviors and for fear conditioning driven by a non-olfactory cue. Our findings identify a selective role for the NLOT in linking olfactory cues to learned aversive outcomes, that supports odor-guided behavioral decisions.

**Significance statement:** Animals rely on smell to evaluate food safety before consuming it, yet the brain circuits that link odor cues to learned aversive outcomes remain poorly understood. Here we identify the nucleus of the lateral olfactory tract (NLOT), a largely unstudied region at the interface between olfactory and limbic circuits, as a critical and selective node for conditioned odor aversion learning. Chemogenetic perturbations of the NLOT disrupted both the acquisition and expression of odor-malaise associations, while leaving odor detection, odor discrimination, and non-olfactory fear learning intact — revealing a previously unrecognized specialization within olfactory-amygdala circuits that supports odor-guided decisions about food safety.

## Introduction

Animals must repeatedly decide whether to consume a specific food source, a choice that depends on both innate preferences and learned experiences (Rozin and Vollmecke, 1986). Consumption of a food followed by aversive post-ingestive consequences, such as malaise or sickness, leads to subsequent avoidance of that food through learned associations between its sensory cues and the negative outcome (Garcia et al., 1955; Garcia and Koelling, 1966; Reilly, 2009). The sense of smell provides rich pre-ingestive information about potential food sources and is therefore a major contributor to consumption decisions (Stutz et al., 2016; Schmitt et al., 2018, 2020; Scherer et al., 2019). However, how olfactory representations are routed to and integrated with outcome representations remains poorly understood. Given the importance of this function, one may predict the existence of dedicated circuits for such associations.

The basolateral amygdala (BLA) is a key structure implicated in forming associations between sensory cues and aversive outcomes (Fanselow and LeDoux, 1999; Maren, 2001), and is part of broader circuits supporting odor-guided learning such as in conditioned odor aversion and taste-potentiated odor aversion paradigms (Hatfield et al., 1992; Miranda et al., 2007; Estrade et al., 2019). Current models suggest that olfactory cortical regions provide detailed representations of odor identity (Gottfried, 2010; Wilson and Sullivan, 2011; Blazing and Franks, 2020), whereas the BLA is broadly implicated in assigning learned significance to sensory cues to guide future behavior (Janak and Tye, 2015). However, the circuit mechanisms linking olfactory representations to the BLA during aversive learning remain poorly understood. In particular, it is unknown where within this circuit odor representations become associated with aversive outcomes through experience-dependent plasticity.

The nucleus of the lateral olfactory tract (NLOT) is a candidate structure for such a role, given its anatomical position within the broader olfactory-amygdala circuit. It receives input from and projects to olfactory regions including the OB and piriform cortex, and also maintains bidirectional connectivity with the BLA (Price, 1973; Broadwell, 1975; Krettek and Price, 1978; Penker et al., 2024), positioning it within circuits linking sensory representations of odor identity and limbic circuits involved in learned odor-guided behavior. The NLOT, however, remains only scarcely studied, and its function is largely unknown. Nonetheless, a small number of studies have hinted at NLOT involvement in olfactory processing, reward-related coding, and transcriptional responses linked to amygdala-associated learning (Vaz et al., 2017; Tanisumi et al., 2021; Hochgerner et al., 2023), though none of these studies have established a causal role for the NLOT in odor-outcome learning.

Together, these anatomical features and the limited but indirect evidence for NLOT involvement in olfactory and learning-related processes position the NLOT as a candidate node within olfactory-amygdala circuits that could contribute to odor-outcome learning. We therefore hypothesized that the NLOT is required for the acquisition and/or expression of learned odor-aversion behavior. Using targeted chemogenetic perturbations across a series of behavioral paradigms, we found that the NLOT activity is required for the acquisition of conditioned odor aversion, but is not necessary for other odor-guided behaviors requiring odor discrimination and detection, or for fear conditioning driven by a non-olfactory cue — indicating a selective role for the NLOT in linking odor cues to aversive outcomes, rather than a general contribution to olfactory processing or associative learning.

## Methods

All procedures were performed using approved protocols in accordance with institutional (Hebrew University Institutional Animal Care and Use Committee) and national guidelines.

### Animals

We used the Rbp4-Cre mouse line (031125-UCD, MMRRC). In these mice the NLOT is the only brain region that both expresses Cre robustly and has axonal projections to the olfactory bulb allowing specific infection of NLOT using cre-dependent retrograde AAVs injected into the OB (Penker et al., 2024). Mice were housed in a reversed light/dark cycle facility and all behavioral training and testing was done during the subjective night time. Both males and females were used.

### Conditioned odor aversion protocol (COA)

The COA protocol was performed in an arena (42 × 26 × 18 cm), equipped with water ports on the two far sides. The water ports were positioned slightly outside the arena requiring mice to protrude their snout through small openings to reach them. At the beginning of each session, water drops were dispensed at both ports and remained available for collection. Water collection was monitored with a capacitance sensor (SparkFun AT42QT1010) and a replacement drop was delivered whenever a drop was collected. Odors were placed in small tubes positioned right underneath the water ports ensuring that mice smelled them while drinking. The COA protocol lasted 4 days, and mice were water restricted starting one day prior to the experiment. On each day, mice were placed in the arena and allowed to drink freely from either side while the number of drops supplied at each port was monitored. On the first day, no odors were presented. On the second day, the non-conditioned odor was positioned under the water spouts on both sides. On the third day, the conditioned odor was positioned on both sides and mice were allowed 10 minutes of drinking. LiCl was then injected intraperitoneally (170 mg/kg, Sigma) to cause gastrointestinal discomfort (malaise). On the fourth day, the conditioned andx non-conditioned odors were positioned on opposite sides. Conditioning was assessed by a preference index PI defined as 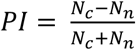, where *N*_*c*_ and *N*_*n*_ are the numbers of drops obtained from the conditioned odor and the neutral odor sides, respectively. In experiments in which NLOT activity was manipulated with DREADDs, Clozapine-N-oxide (CNO, 10 mg/kg) was injected intraperitoneally 2 hours prior to the behavioral session. The odorants used were Isoamyl Acetate (>95%, cas 123-92-2, Sigma-Aldrich) and Hexenal (98%, cas 66-25-1, Sigma-Aldrich) and each odorant served as the conditioned odor for some mice and as the non-conditioned odor for others.

Statistical testing of conditioning results was performed by fitting a linear mixed-effects model (Matlab fitlme). The model was specified as preference *∼ group × day + (day* | *mouse)*. Overall fixed effects were assessed using an ANOVA on the fitted model. To evaluate between-group differences in learning, custom contrast vectors were used to compare the interaction terms of each experimental group against the control group (Matlab coefTest). Additionally, within-group learning was assessed for each group separately using contrasts that isolated the effect of day for each group by combining the main effect of day with the relevant group-by-day interaction term. All resulting p-values were corrected for multiple comparisons using the Benjamini-Hochberg false discovery rate (FDR) procedure, applied separately to the between-group contrasts and the within-group learning contrasts (Benjamini and Hochberg, 1995).

### Auditory fear conditioning

Fear conditioning was conducted in a chamber with a grid floor connected to a shock generator and enclosed within a sound-attenuating acoustic chamber. Mice were placed in the chamber for 2 min, after which a pure tone (4 kHz, 20 s) was presented and co-terminated with a 2 s foot shock (0.6 mA); this pairing was repeated once, and mice were removed 30 s after the second shock. Fear responses were quantified by continuous measurement of freezing behavior, defined as complete immobility except for respiration. Contextual fear memory was assessed on day 2 by returning mice to the original conditioning chamber and measuring freezing for 5 min, while cued fear memory was tested by placing mice in a novel context with altered walls and floor for 2.5 min, followed by tone presentation for an additional 2.5 min during which freezing was measured.

Statistical analysis of conditioning results was performed by fitting linear mixed-effects models as detailed above for odor conditioning.

### Urine engagement

In this experiment, only males were tested and female urine was used for monitoring engagement. Urine was collected on the day of the experiment from 5-6 female mice. The urine was smeared on a standard mouse cage’s wall at one random corner. Mice were placed individually in the center of the cage and their location was video recorded for 5 min. Attraction to the urine was quantified for each mouse by comparing the distribution of the mouse’s distance to the urine corner to the distribution of the mouse’s distance to the opposite corner. These distances were measured for each frame and the difference between the distributions was quantified by the area under the ROC curve. ANOVA was then performed on the obtained values to compare between mouse groups. P-values for attraction for each group were obtained using the Wilcoxon signed rank test and were corrected for multiple comparisons using the Benjamini-Hochberg false discovery rate.

### Buried food test

Animals were food-deprived one day prior to experiment. The chamber consisted of a clean standard plastic cage (42 × 26 × 18 cm) with a 4 cm layer of new bedding. A food pellet (standard mouse chow) was buried in the bedding, approximately 3 cm beneath the surface, at a random location. The bedding surface was smoothed out and the mouse was introduced into the cage. The time until the buried food was found was recorded. The maximum test time allowed was 480s. Statistical analysis of these data were performed using the Kruskal-Wallis test.

### Target-Odor Detection Tasks

#### Surgery

Mice were anesthetized (Isoflurane), the skull was exposed and a metal head-plate was attached with dental adhesive (Metabond) for subsequent head restraining. Viruses were injected as described below. Mice were allowed to recover for 3 week before water restriction and behavioral training.

#### Task

Odor detection and identification tasks were performed by mice expressing DREADDs. CNO was injected before a subset of sessions (2 hour prior to the behavioral session) and performance of individual mice was compared across these two conditions. The task was arranged in a go / no-go paradigm and performed during head-restraint with water-restriction and water rewards for motivation (Rokni et al., 2014; Lebovich et al., 2021; Licht et al., 2023). Every mouse was arbitrarily assigned a target odor. In each trial (every 12 seconds), subjects were presented with a pseudorandom combination of odorants (from a pool of 8 odorants) and were required to lick the water spout if the mixture contained the target odorant (go trial) and refrain from licking if it didn’t (no-go trial). Correct licks were rewarded with a 3 μL drop of water. For some of the mice we then replaced the target odor daily to test their ability to learn new reward contingencies. In this task, only single odors were presented instead of mixtures.

### Virus injections

Mice were anesthetized with isoflurane, administered analgesia (Meloxicam 5 mg/kg), and secured in a stereotactic device (David Kopf Instruments). A small craniotomy was made over the olfactory bulb, and viral solution was injected into the brain with a WPI 10µl syringe (volume, 200 nl at a rate of 100 nl/min). The needle was then slowly retracted, the craniotomy was sealed with bone wax, and the skin was sutured. A minimum of 3 weeks were allowed for all adeno-associated viruses (AAVs) to express. Retrograde AAVs were made at the ELSC Vector Core Facility at the Hebrew University using the following plasmids: pAAV-EF1a-DIO-hM3D(Gq)-mCherry (Addgene Plasmid #50460) for chemogenetic activation and pAAV-hSyn-DIO-hM4D(Gi)-mCherry (Plasmid #44362) for chemogenetic inhibition.

### Histology, immunostaining and microscopy

The expression of DREADDs in all virus-injected mice was confirmed histologically. Brains were fixed by immersion in 4% parafor-maldehyde on ice for 12h, washed with PBS, and sectioned to 50 μm coronal sections by a vibratome (Leica). Staining was performed as described (Licht et al., 2023; Penker et al., 2024) with rabbit anti-cFos antibody (1:600, synaptic systems RRID:AB_2231974). Alexa Fluor 647 anti-rabbit (1:400, Jackson ImmunoResearch Laboratories; RRID, AB_ 2340625) was used as secondary antibody. Sections were mounted on glass slides, covered by mounting medium containing DAPI (SouthernBiotech) and coverslipped. Low-magnification images were acquired using Nikon SMZ-25 fluorescent stereoscope equipped with 1× and 2× objectives. High-resolution images were acquired with an Olympus FV-1000 confocal microscope equipped with 10× and 20× objectives. Images were taken at 3 µm distance between confocal z-slices.

### cFos quantification

cFos expression was quantified by comparing the density of cFos labeling within a defined region of interest (ROI) that of the surrounding using a custom-written Matlab script. ROIs were manually delineated as polygons on 10X confocal images using the DAPI and mCherry channels for anatomical guidance. Mean pixel intensity in the cFos channel was then calculated separately for pixels inside the ROI and for pixels outside the ROI. For each section, a density ratio (ROI / surrounding) was computed, and the distribution of these ratios was compared across experimental groups.

## Results

To test the involvement of the NLOT in various behaviors, we perturbed NLOT activity with Designer Receptors Exclusively Activated by Designer Drugs (DREADDs) and compared mouse behavior to unperturbed controls (Roth, 2016). The small size and ventral location of the NLOT make direct stereotaxic injections technically challenging. To overcome this limitation, we used a genetic targeting strategy that enables precise and highly specific labeling of NLOT neurons. We previously showed that injection of Cre-dependent retrograde AAVs into the main olfactory bulbs (MOB) of Rbp4-cre mice leads to specific and extensive expression of targeted genes in layer 2 of the NLOT (Penker et al., 2024). We therefore injected Cre-dependent retrograde AAV particles carrying the gene for either the inhibitory hM4Di or the excitatory hM3Dq bi-laterally into the MOBs of Rbp4-cre mice (Fig 1A, B). We first tested the efficacy of the DREADD manipulations by fixing mouse brains 2 hours after intraperitoneal administration of the DREADD agonist Clozapine-N-oxide (CNO) and immunostaining for the immediate early gene c-Fos. Mice expressing the excitatory DREADD hM3Dq showed a 25-fold increase in c-Fos staining specifically in the NLOT following CNO administration indicating successful activation of these neurons (Fig 1C, D). As the NLOT projects back to the OB bilaterally, even unilateral injections yielded bilateral expression and elevation of cFos levels, although cFos levels in the contralateral NLOT were not as prominent as the ipsilateral NLOT. No reduction of c-Fos immunostaining was evident in mice expressing the inhibitory hM4Di, presumably due to the very low levels of c-Fos expression in controls (Fig 1C, D).

**Figure 1:**
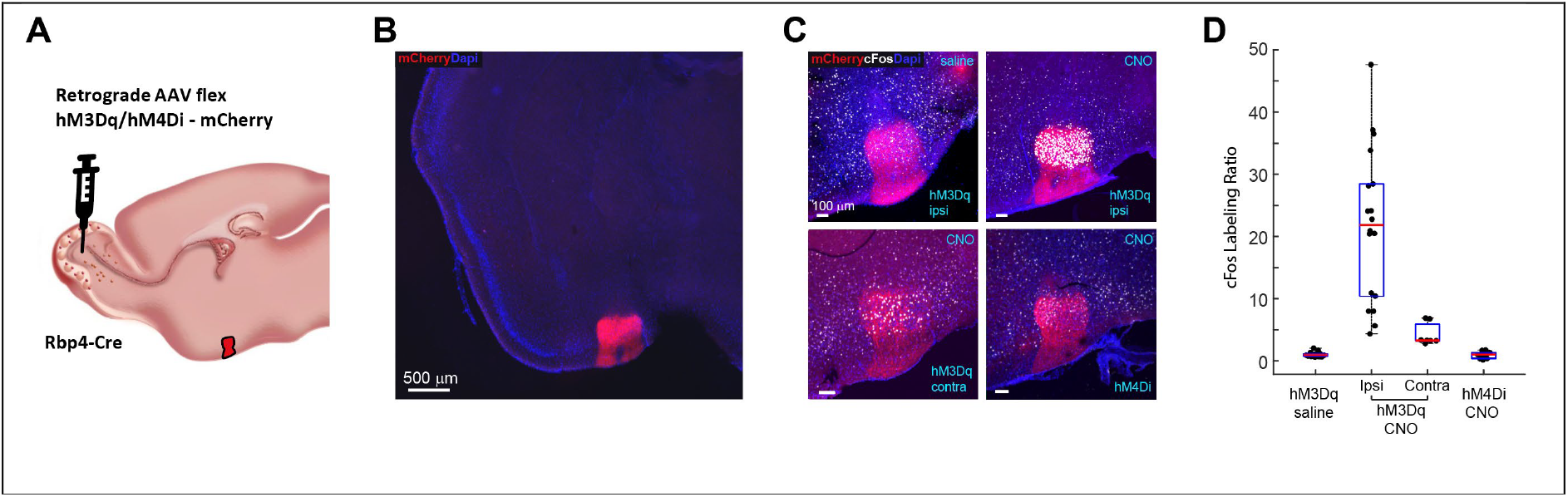
NLOT-specific manipulation of neuronal activity with DREADDs. **A**. Schematic of infection of the NLOT with DREADD-encoding retrograde AAVs injected into the MOB. To assess contralateral effects, we injected the virus unilaterally. **B**. A coronal section showing the specific expression of cre-dependent AAVs in the NLOT of Rbp4-Cre mice using the fluorescent reporter mCherry. **C**. Immunostaining for cFos in brains that were fixed 2 hours after i.p. administration of either CNO or saline in mice expressing DREADDs. **D**. Quantification of cFos labeling 2 hours post saline/CNO injection in mice expressing DREADDs. The cFos labeling density inside the NLOT was normalized by labeling density outside the NLOT to obtain the labeling ratio in each section. Kruskal-Wallis P = 10^-8^ effect of mouse group, post hoc comparisons: p = 10^-7^ hM3Dq ipsi vs saline, p = 0.04 hM3Dq contra vs saline, p = 1 hM4Di vs saline (n=18, 17, 7, and 10, for hM3Dq saline, hM3Dq ipsi, hM3Dq contra, and hM4Di, respectively).

We examined how manipulating NLOT neuronal activity affects both the learning and the expression of conditioned odor aversion (COA). In this paradigm, mice learn to associate a specific odor presented during water consumption with subsequent malaise, leading them to later avoid drinking water in the presence of that odor. In this four-day behavioral protocol, water-restricted mice are placed in an arena containing two opposing water ports and allowed to drink freely from either side (Fig 2A, B). After an acclimatization day (Day 1), odor sources are introduced near the ports. On Day 2, a neutral non-conditioned odor is positioned under both water ports. On Day 3 – the conditioning session -the conditioned odor is positioned under both ports and mice are allowed 10 minutes of free drinking, after which they receive an intraperitoneal injection of lithium-chloride (LiCl) to induce malaise. On the test day (Day 4), the conditioned and non-conditioned odors are placed on opposite sides, and the drinking preference of the mice serve as the measure of conditioned aversion.

**Figure 2:**
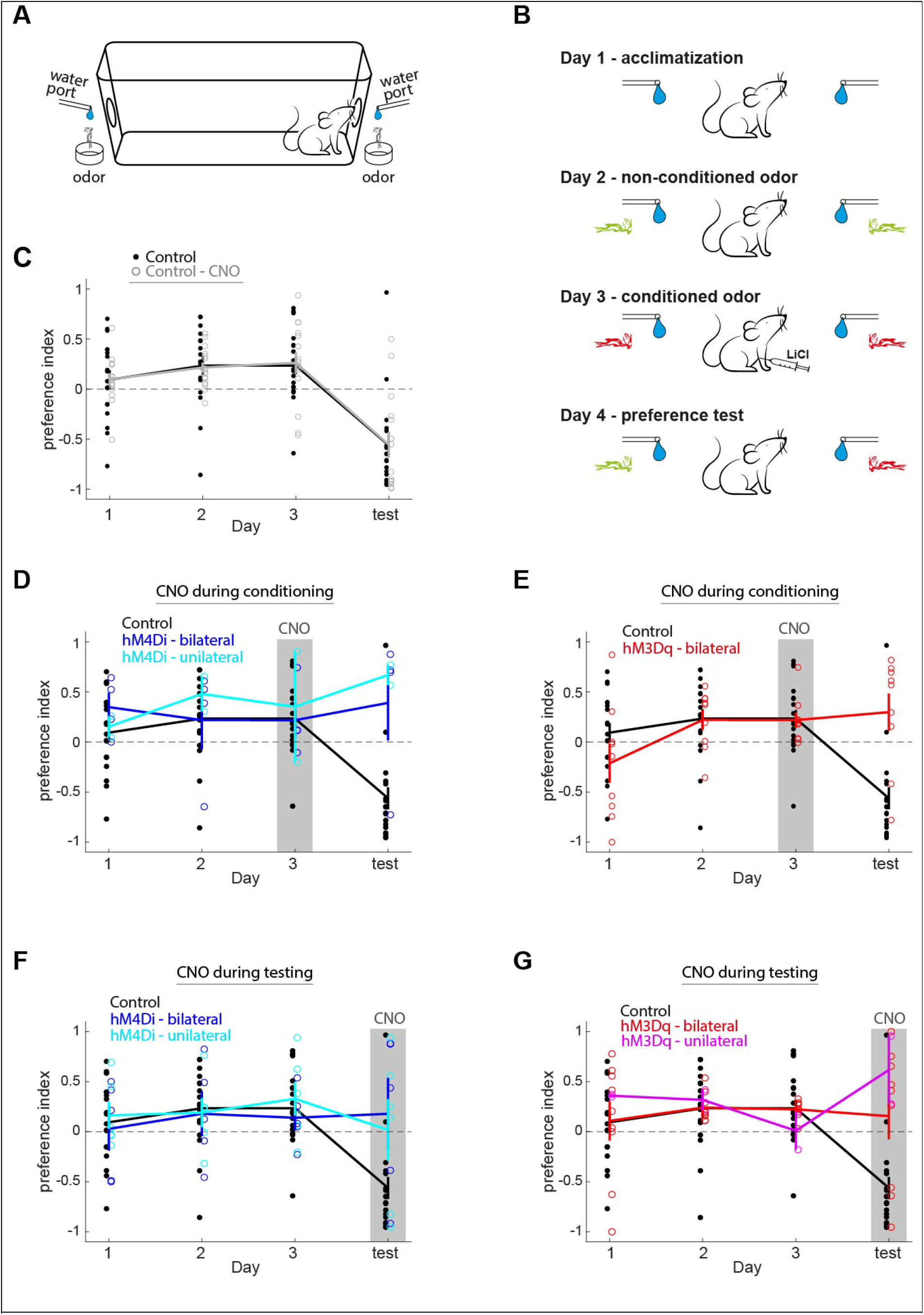
Effects of NLOT perturbations on COA. **A**. A schematic of the conditioning arena. **B**. The conditioning protocol. **C**. Drinking preferences of untreated mice (black, n=18), and CNO-administered control mice (gray, n=15) along the days of the conditioning protocol. Positive and negative values indicate preference and avoidance of the conditioned-odor side, respectively (see Methods). Both groups show significant conditioning (p < 0.001 for both groups), with no significant difference between them (p = 0.8). **D and E**. Conditioning results in mice expressing the inhibitory hM4Di (D, n=6) and excitatory hM3Dq (E, n=9) and administered with CNO two hours prior to the conditioning session. Both groups show significant differences from both control groups (p = 0.0005 vs non-injected controls for both D and E, and p = 0.0002 vs CNO-injected controls, for both D and E), with neither group showing significant learning (p = 0.47 and 0.68, for D and E, respectively). Blue and cyan in E indicate bilateral and unilateral injection of the virus, respectively. **F and G**. Same as D and E but with CNO administered two hours prior to the test session (n=9 and 11 for F and G, respectively). Here too the NLOT manipulation prevented the expression of conditioned aversion (p = 0.012 and 0.002 for comparison to non-injected controls, and p = 0.005 and 0.0006 vs CNO-injected controls, for F and G, respectively; p = 0.56 and 0.77 for comparisons between Day 3 and test day for F and G, respectively). For D-G, blue and red indicate bilateral injection of the virus, while cyan and magenta indicate unilateral injection of the virus. P values for all conditioning experiments were obtained by analyzing fitted linear mixed-effects models (see Methods). Bilateral and unilateral expression were combined for this analysis. All reported p-values were corrected for multiple comparisons using the Benjamini Hochberg false discovery rate procedure.

Control mice show clear avoidance in the test session, indicating successful learning of the odor-malaise association (Fig 2C). To test whether the NLOT is part of the circuitry underlying the formation of this association, we compared conditioning in control mice and during chemogenetic manipulation of NLOT neuronal activity with DREADDs. We first tested whether the DREADD agonist CNO, has any non-specific effects on COA. To this end a group of mice that were not infected with DREADDs was injected with CNO two hours prior to the conditioning session. These mice showed clear avoidance during the test session, no different than that of the controls indicating that CNO by itself has no effect on COA (Fig 2C).

Next, we tested whether inhibition of the NLOT interferes with COA. To this end we infected mice with hM4Di and administered CNO two hours before the conditioning session. No conditioned preference was evident during the test session in these mice (Fig 2D). This effect was also evident in mice in which virus injection into one of the hemispheres failed, possibly reflecting the bilateral NLOT to MOB projections, as well as the interhemispheric interactions between the two NLOTs (Penker et al., 2024). To test the effect of NLOT activation on COA, we expressed the excitatory DREADD hM3Dq in the NLOT of another group of mice. CNO administration prior to conditioning in these mice completely abolished avoidance on the test day, similarly to the inhibitory DREADD (Fig 2E). We next tested whether the NLOT is also required for retrieval of the association. To this end, we injected CNO two hours before the test session while leaving NLOT activity intact during the conditioning session (Fig 2F and G, respectively). These mice also failed to exhibit conditioned aversion, indicating that perturbation of the NLOT blocks the expression of learned COA. These data indicate that intact NLOT activity is required for COA and that perturbations of its neuronal activity block both learning and expression of COA, and that this effect is irrespective of perturbation directionality (inhibition or activation).

The results presented so far suggest that the NLOT serves a function of associating odors with the malaise, however there two alternative interpretations. First, as the NLOT is part of the olfactory circuitry and is interconnected with olfactory regions, manipulating its activity may interfere with olfactory processing and may affect the detection or the identification of the conditioned odor. Second, as the NLOT is also interconnected with the BLA, DREADD manipulations of NLOT activity may result in perturbation of BLA activity and potentially interfere with learning any negative valence association irrespective of modality.

To test whether manipulating NLOT activity perturbs odor perception, we tested mice on pure olfactory tasks that test various aspects of olfactory function. We started with simple self-motivated tasks and then trained mice on structured tasks that directly inform of odor identification capabilities. We started with the buried food test in which a food pellet is placed underneath the cage bedding and hungry mice are allowed to roam and find it. The time until they find the food is used as a crude measure of olfactory function. We measured this time in control mice as well as mice expressing either hM3Dq or hM4Di. The DREADD-expressing mice were injected with CNO two hours prior to behavioral testing. All mouse groups found the food with a median of around 1 minute, with no statistically significant differences between the groups (Fig 3A). We next tested whether NLOT manipulations affect how mice interact with a naturally meaningful smell – opposite sex urine. To this end, we placed a few drops of female urine in one corner of a test cage and introduced male mice into the cage for 5 minutes. Mice were allowed to roam freely and were continuously monitored. The amount of time they spent in vicinity to the urine was taken as a measure of odor-guided interest. All mouse groups showed a significant preference towards the corner with the urine, and here too we found no significant difference between the groups (Fig 3 B, C). Taken together, these experiments show that the same perturbations that affected COA, did not affect self-motivated odor-guided behaviors.

**Figure 3:**
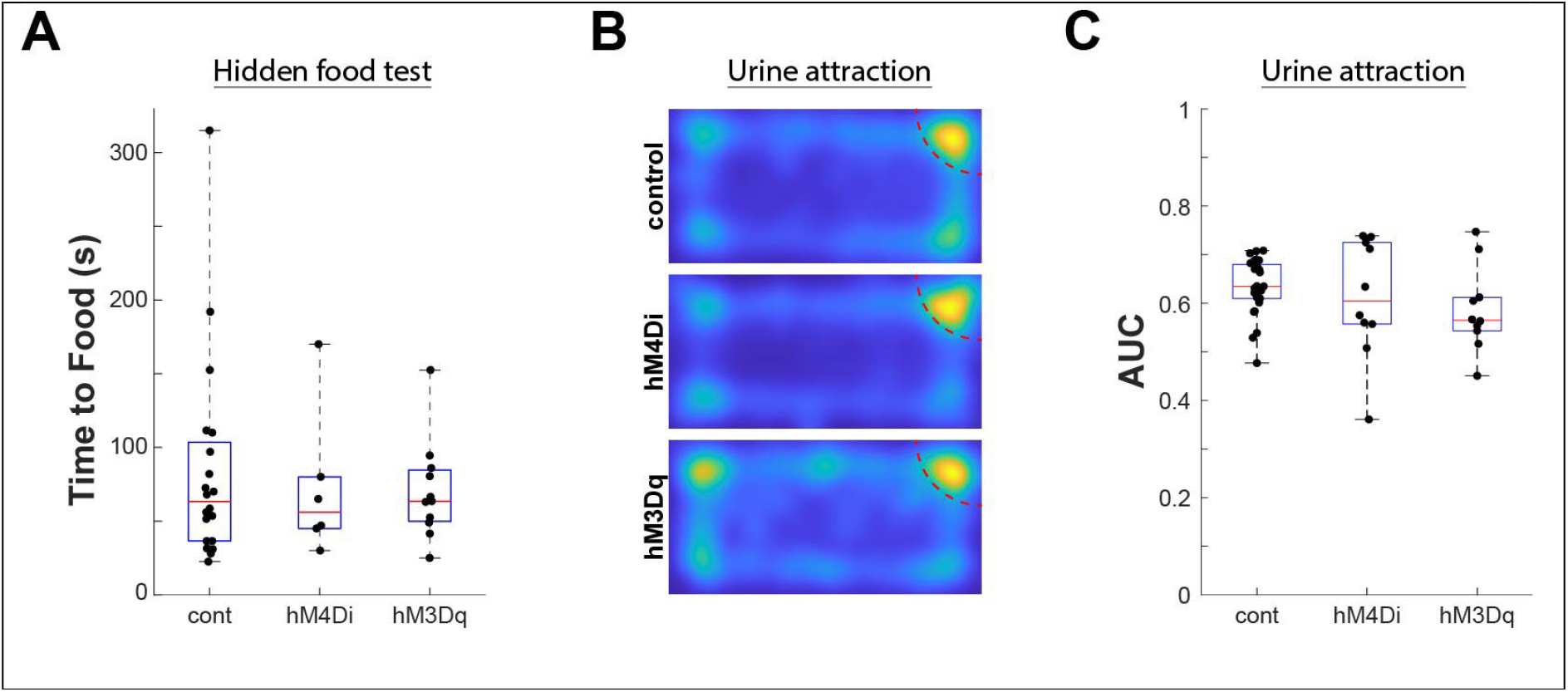
Effects of NLOT perturbations on self-motivated odor-guided behaviors. **A**. The time it took for mice to find food that was buried under the cage bedding. Mice were food-deprived for 24 hours prior to the test. A Kruskal-Wallis test revealed no effect of group (p = 0.12, n = 20, 6, and 12, for controls, hM4Di, and hM3Dq, respectively). **B**. Place preferences of male mice that were introduced into a cage in which female urine was presented in the top right corner. The heat maps show the distributions of time spent in each location in the cage, averaged across mice within each group. **C**. Quantification of place preference. For each mouse, the distribution of its distances to the urine corner was compared to the distribution of its distances to the opposing corner using ROC analysis (see methods). A Kruskal-Wallis test revealed no significant differences between the groups (p = 0.15, n = 18, 10, and 10, for controls, hM4Di, and hM3Dq, respectively), with hM3Dq being the only group that was significantly different than controls (p = 0.017 post hoc analysis).

These self-motivated behaviors do not directly test odor identification abilities. We therefore next turned to a structured task in which mice are trained to report odor identification. Specifically, we used an olfactory figure-background segmentation task (Rokni et al., 2014; Lebovich et al., 2021). In this task, mice are presented with mixtures of odorants and are required to report for each, whether it contains a particular target odorant. Mice report detecting the target odorant by licking a water port in front them and receive a water reward for correct licks. Performing this task requires target odor identification that is further made difficult by odor distractors and therefore provides a good measure for odor acuity. We trained mice expressing either hM3Dq or hM4Di and tested the effect of CNO on task performance by interleaving control sessions and CNO sessions and comparing performance between these cases directly. We found that CNO did not alter the overall performance in this task in either mouse group (Fig 4A). We further analyzed whether CNO had any specific effect on performance in trials with more distractors. Figure 4B shows very similar performance with and without CNO irrespective of the number of mixture components. There was no statistically significant difference between session with and without CNO in trials with 6 or 7 mixture components. These data indicate that a smell that is associated with a specific meaning without CNO (here with the requirement to lick and obtain a reward) can still be identified when the NLOT is perturbed with DREADDs. We conclude that the NLOT is not required for odor identification and rule out the possibility that the effects of NLOT perturbations on COA were due to effects on odor perception.

**Figure 4:**
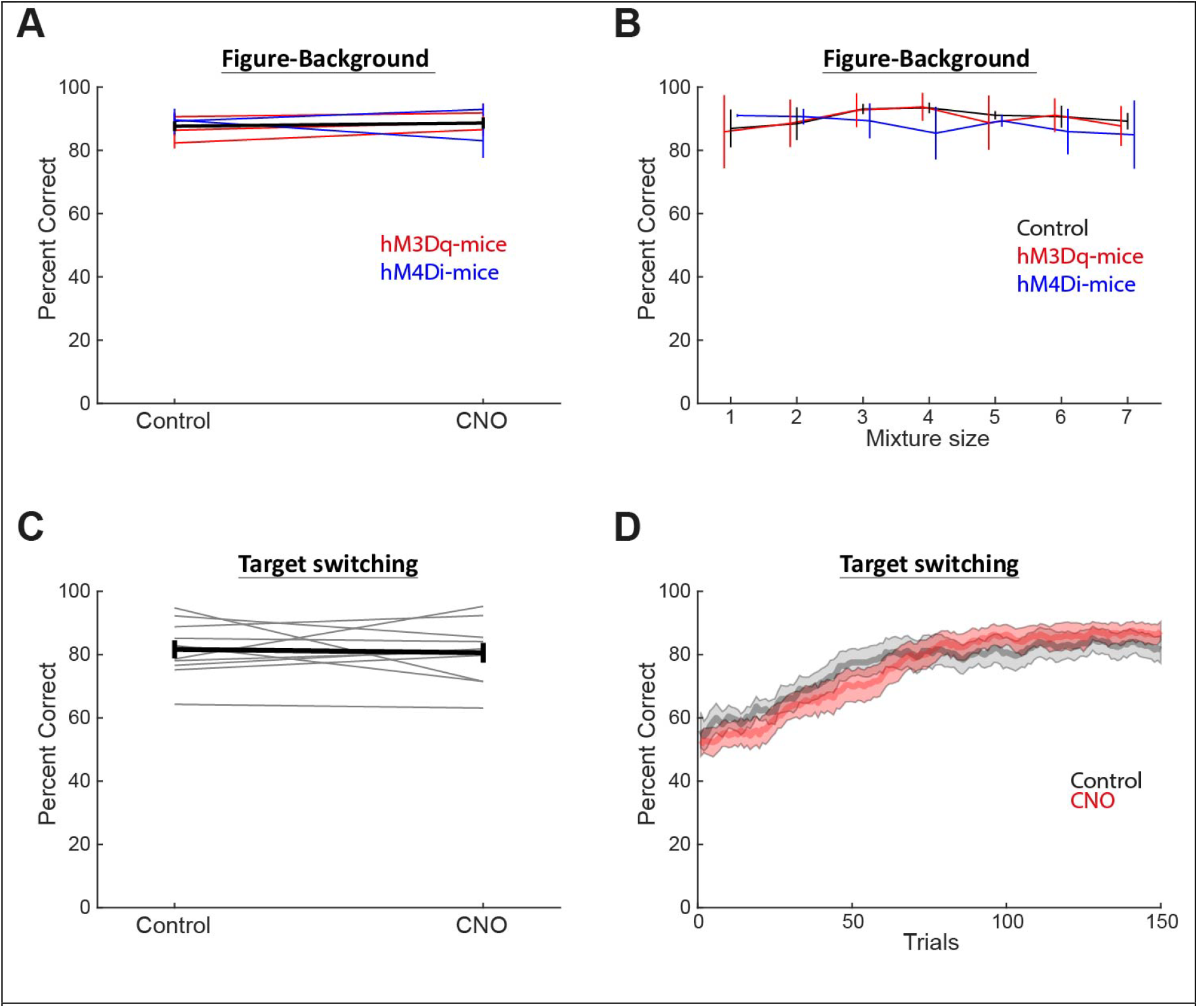
Effects of NLOT perturbations on odor identification tasks. **A**. Performance of mice expressing DREADDs on an olfactory figure-background segmentation task with and without CNO administration. Blue and red lines show the average performance of hM4Di and hM3Dq mice, respectively (N = 2, and 3 mice, between 450 and 2200 trials each). Black line shows the averages of all mice. **B**. The average performance of each mouse group as a function of mixture size. Blue, and red lines show data from mice expressing hM4Di and hM3Dq, respectively (N = 2, and 3 mice, between 450 and 2200 trials each) and administered with CNO. Black line shows data from the same mice in control sessions. **C**. Performance of mice expressing hM4Di on a Go-NoGo task with daily switching of the target-odorant, in control and CNO sessions. Each gray line shows two consecutive sessions of which the first was a control session and the second a CNO session (10 session-pairs from 2 mice). Black line shows the average of all session in both conditions. **D**. A running average (25 trials window) of the performance as a function of trial number for the sessions in C. Shaded areas show SEM. Black shows control sessions and red shows CNO sessions.

To test whether NLOT perturbations abolish the ability to learn any odor associations, we tested another group of mice, expressing hM4Di, on a paradigm in which the target odor changes every day. Here, mice needed to learn a novel association every day and we compared this ability with and without CNO administration. To alleviate a potential credit assignment problem, only single odorants were presented in each trial rather than mixtures as before. Here too, CNO had no effect on the overall performance in this task (Fig 4C). The learning speed within individual sessions was very similar between CNO and control sessions as well (Fig 4D). These data indicate that the NLOT is not required for assigning significance to odors in this form of appetitive learning.

As mentioned earlier, NLOT neurons also communicate bidirectionally with the BLA which is presumed to be important for most aversive conditioning paradigms, including COA. Thus, manipulating NLOT activity may disrupt BLA function in a modality-nonspecific fashion. Particularly, NLOT neurons have been shown to project to inhibitory interneurons within the BLA so their activation with DREADDs might inhibit BLA projection neurons and thereby block conditioning (Krabbe et al., 2019). To test this, we first tested whether DREADDs affect cFos expression in the BLA. We found that neither activation of hM3Dq nor inhibition via hM4Di affected cFos expression in the BLA (Fig 5A, B). We next tested the effect of NLOT manipulations on the acquisition of auditory fear conditioning, in which a benign sound is associated with an electrical foot shock. Both control mice, and mice expressing either hM3Dq or hM4Di and administered with CNO, demonstrated increased freezing in response to the tone in the retrieval test, with no significant difference between each manipulation and the control group (Figure 5C, D). We therefore rule out the possibility that NLOT perturbations caused a general modality-unspecific perturbation of BLA activity and conclude that the NLOT plays a specific role in linking olfactory stimuli with negative outcomes.

**Figure 5:**
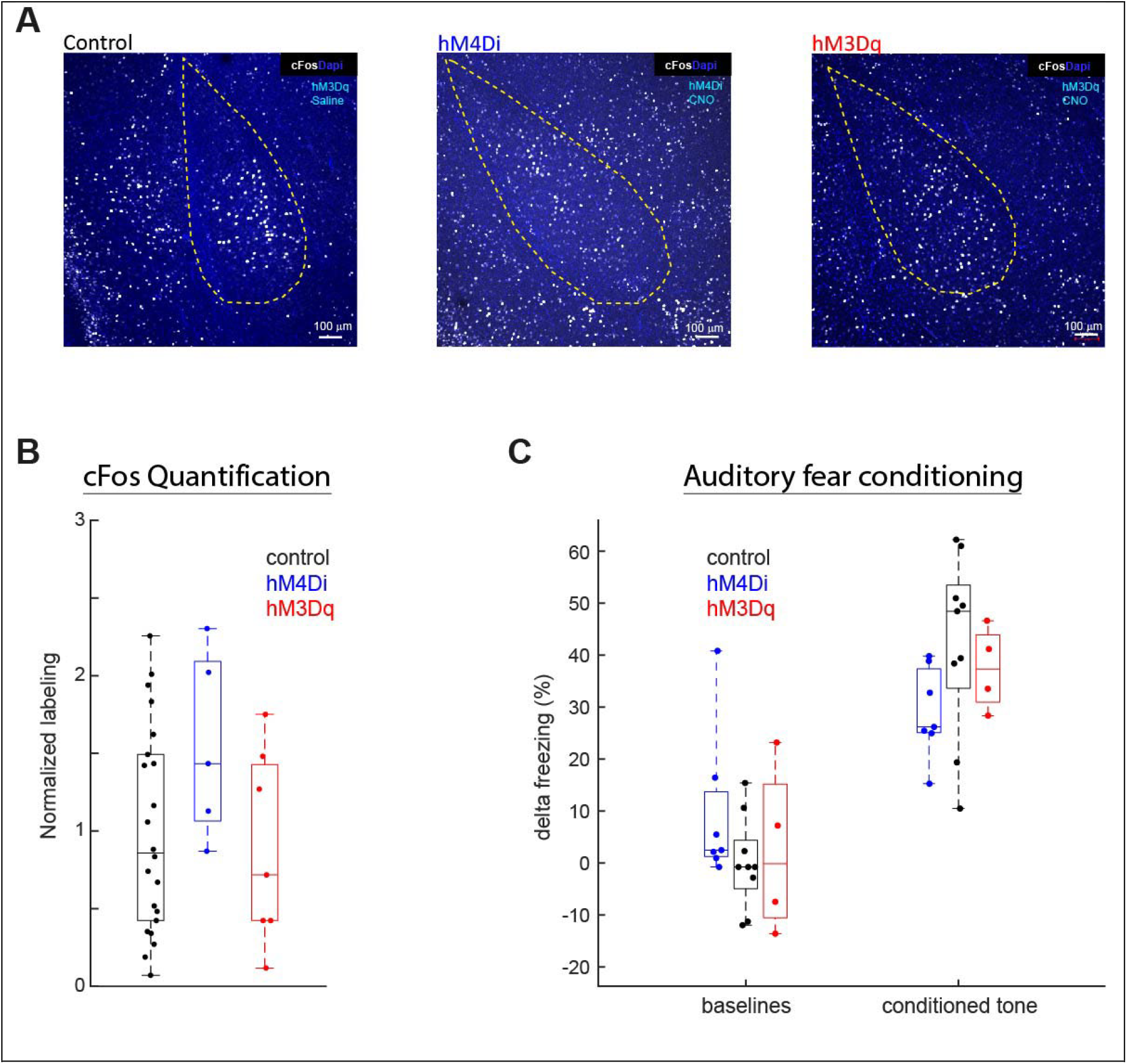
Effects of NLOT manipulations on acquisition of auditory fear conditioning. **A**. Example images showing immunolabeling of cFos in the BLA in mice expressing DREADDs. **B**. The ratio of labeling density inside and in the vicinity of the BLA. P=0.18, Kruskal Wallis test, n = 22,5, and 7, for control, hM4Di, and hM3Dq, respectively. **C**. Freezing behavior before and after the conditioning paradigm in control mice and mice expressing either hM4Di or hM3Dq. The average baseline freezing of all mice in a session was subtracted for each mouse. Analysis of a fitted linear mixed effects model revealed that each of the three groups showed a significant increase in freezing (p = 10^-5^, 0.001, and 0.02, for controls, hM4Di, and hM3Dq, respectively), with no significant difference between either of the DREADD groups and controls (p=0.1 and 0.15 for hM4Di, and hM3Dq, respectively).

## Discussion

We combined chemogenetic manipulations with behavioral assays to define the functional role of the NLOT and found that it is required for conditioned odor aversion. NLOT perturbations impaired acquisition of conditioned odor aversion (COA), but did not affect odor detection or discrimination, indicating that basic perception of odor identity remained intact. In addition, NLOT manipulations did not affect auditory fear conditioning, suggesting that its function is not required for associative fear learning in a non-olfactory modality. Together, these findings indicate that NLOT activity is specifically required for conditioned odor aversion learning rather than for odor perception or general fear/aversive learning.

Chemogenetic perturbations using both either the excitatory hM3Dq and inhibitory hM4Di yielded similar behavioral effects, suggesting that normal NLOT function depends on appropriate patterns of activity rather than simple increases or decreases in overall excitability. Both over-activation, and inhibition likely perturb these patterns and therefore interfere with NLOT function. At the circuit level, the net influence of NLOT input onto basolateral amygdala output neurons remains incompletely understood. Anatomical tracing indicates that NLOT inputs target multiple inhibitory cell types within the BLA, including both interneuron classes that suppress and disinhibit principal neurons. Whether NLOT inputs also directly target BLA projection neurons remains to be determined, and will be important for understanding how NLOT activity shapes downstream circuit computations.

What role the NLOT plays in conditioned odor aversion remains unclear. One possibility is that the NLOT itself participates directly in odor-outcome learning by integrating odor representations with aversive teaching signals within the olfactory-amygdala circuit. Alternatively, the NLOT may function primarily as an intermediate node that shapes odor-related input to downstream structures such as the BLA, where associative plasticity could occur. These possibilities are not mutually exclusive, and both are consistent with the observed requirement of NLOT activity for COA. Given its reciprocal connectivity with both olfactory regions and the BLA, the NLOT is well positioned to influence both sensory representations and downstream limbic processing, making either mechanism plausible. More broadly, our findings establish the NLOT as a necessary component of the circuit supporting conditioned odor aversion, without resolving the locus of learning-related plasticity.

Several features of the NLOT make the possibility of local plasticity an interesting hypothesis for future investigation. First, the NLOT receives dense projections from cholinergic neurons (Millhouse and Uemura-Sumi, 1985; Penker et al., 2024), inputs that have been implicated in modulating synaptic plasticity during associative learning (Letzkus et al., 2011; Jiang et al., 2016; Crimmins et al., 2023). While the functional role of these inputs in the NLOT remains unknown, they provide a potential mechanism for learning-related modulation of its function. In addition, NLOT neurons exhibit burst firing and calcium-dependent spiking dynamics that have been shown to promote plasticity in other systems (Pike et al., 1999; Bittner et al., 2015, 2017). Together with the behavioral requirement for NLOT activity demonstrated here, these properties suggest that the NLOT may be well positioned for local experience-dependent modulation, raising the possibility that learning-related changes could occur within the NLOT itself. Future experiments will be required to directly test whether and how NLOT plasticity contributes to odor conditioning.

Whether the NLOT is generally required for aversive odor learning across paradigms, or instead plays a more selective role in ingestion-associated conditioned odor aversion, remains unclear. Odor-foot shock fear conditioning, the most extensively studied olfactory aversive learning paradigm, depends on interactions between olfactory cortical regions and the basolateral amygdala and is associated with experience-dependent changes across multiple nodes in the olfactory system (Sevelinges et al., 2004; Kass et al., 2013; Gore et al., 2015; Fast and McGann, 2017; Ross and Fletcher, 2018; Meissner-Bernard et al., 2019; East et al., 2021; Bakir et al., 2026). However, the involvement of the NLOT in this form of learning has not been examined. Notably, odor-shock fear conditioning differs from COA in its temporal structure. Whereas fear conditioning pairs odors with an immediate aversive event, COA associates odors with malaise that develops minutes after consumption, which is thought to require specialized neural mechanisms (Adaikkan and Rosenblum, 2015; Zimmerman et al., 2025). Whether the NLOT contributes broadly to aversive odor learning or is specifically important for these temporally extended odor-outcome associations remains an open question.

We find a role for the NLOT in learning aversive value. Whether it may also play a role in positive valence associations is not clear, yet our result that mice can relearn the target odor in an odor-identification task during NLOT perturbations, suggests that the NLOT is not required for all forms of reward-guided odor behavior. Previous studies suggested that assignment of positive value to odors occurs in the olfactory striatum (Gadziola et al., 2015; Millman and Murthy, 2020). Together, these findings are consistent with the idea that distinct downstream olfactory circuits contribute differentially to odor-guided behavior, potentially supporting parallel routes through which odor representations can acquire learned significance. Importantly, however, the present data do not distinguish whether these pathways reflect specialized “valence modules” or more general learning circuits engaged by different task demands.

In summary, our results identify the NLOT as a critical and previously unrecognized node in the circuitry required for linking odor representations to delayed ingestion-related aversive outcomes. Rather than serving a general role in olfactory processing or aversive learning, the NLOT appears to be selectively required for this form of odor-outcome learning, highlighting a previously unappreciated level of specialization within olfactory-amygdala circuits. These findings identify the NLOT as a key interface between olfactory and learning-related circuits and establish a foundation for future studies to determine whether its role extends to other forms of odor-guided aversive learning and to identify the circuit mechanisms by which odor representations become associated with negative outcomes.

## Acknowledgements

This study was funded by an Israel Science Foundation grant (1188/23)

